# Constructing Graphs from Genetic Encodings

**DOI:** 10.1101/2020.11.02.365189

**Authors:** Dániel L. Barabási, Dániel Czégel

## Abstract

Our understanding of real-world connected systems has benefited from studying their evolution, from random wirings and rewirings to growth-dependent topologies. Long overlooked in this search has been the role of the innate: networks that connect based on identity-dependent compatibility rules. Inspired by the genetic principles that guide brain connectivity, we derive a network encoding process that can utilize wiring rules to reproducibly generate specific topologies. To illustrate the representational power of this approach, we propose stochastic and deterministic processes for generating a wide range of network topologies. Specifically, we detail network heuristics that generate structured graphs, such as feed-forward and hierarchical networks. In addition, we characterize a Random Genetic (RG) family of networks, which, like Erdős-Rényi graphs, display critical phase transitions, however their modular underpinnings lead to markedly different behaviors under targeted attacks. The proposed framework provides a relevant null-model for social and biological systems, where diverse metrics of identity underpin a node’s preferred connectivity.

## Introduction

Generative models in network science are often designed to recreate unique topologies observed in nature [1, 2, 3, 4]. Such approaches range from the random edge insertion of the Erdős-Rényi model [5] to the preferential attachment model [6], which aims to capture the growth of certain real-world networks. Although these models utilize interpretable processes, they each produce a family of networks with specific properties, such that ER networks are rarely scale-free, and preferential attachment leads to highly non-random topologies. On the other hand, statistical methods, like the maximum entropy approach, can generate graphs with custom properties [7, 8], but the utilized sampling processes are not expected to match the evolution of real systems.

Recent work in network neuroscience has elucidated how the reproducible connectivity of a brain can arise from known developmental processes and quantifiable genetic interactions [9]. Specifically, neural connectivity is encoded in part by genetic compatibility: if genes A and B are compatible, then all neurons expressing gene A have the potential to link with all neurons expressing gene B. Thus, a direct mapping can be found between a neuron’s (node’s) identity and its connectivity [10]. This can be seen as a specific network encoding method: rather than producing a family of networks, a known set of genetic interactions reproducibly generates a single graph.

Drawing inspiration from the physical laws apparent in brain connectivity, we propose using genetic encoding as a general way to construct a network, thereby defining a generative model that results in topologies built out of a set of modular components. To show the model’s utility, we define network encodings that use simple heuristics to generate scale-free and feed-forward topologies, highlighting the flexibility of this representation. We then propose a Random Genetic (RG) model, which resembles the ER model, but samples wiring rules instead of edges. We find that the RG model generates a novel family of networks with (1) a characteristic degree distribution and (2) a connectivity structure with three phases, corresponding to subcritical, critical, and supercritical link density. Our work elucidates how genetic encoding can balance the generation of structured graphs with the exploration of new topologies through stochastic processes.

### Encoding Known Topologies with Genetic Rules

To illustrate the genetic encoding of a network, we show how wiring rules can provide the basis for two well-known and highly dissimilar architectures: a scale-free and a feed-forward network. We begin with the two principles of the Connectome Model [9]:

First, we assume that nodes are uniquely labeled by a binary barcode. To be specific, we consider nodes to be uniquely labeled by the combinatorial expression of *b* bits. For instance, given b = 2, we can have nodes with barcodes 00, 01, 10, and 11. This allows for a total of 2^*b*^ unique nodes over the identity space {0, 1}^*b*^.

Second, we consider connections to be introduced through “wiring rules” that recognize and connect subsets of nodes based on shared bits, thereby forming bicliques [9]. Indeed, consider a system with *b* = 5, where 2^*b*^ = 32 nodes exist in this system. For the moment we focus on only 8, which have been divided in Sets A and B (Figure 1a, top). Set A contains all nodes that have bits 1 and 3 on, but have bit 2 off (Figure 1a, left). We use the shorthand 101XX to express the membership of Set A, where Xs stand for “any” bit state, thus nodes with barcodes 10100, 10101, 10110, and 10111 all belong to Set A. In this formalism, Set B is defined as 110XX, or all nodes with bits 1 and 2 on, but bit 3 off (barcodes: 11000, 11001, 11010, and 11011). We now observe that all nodes in set A connect to all nodes in set B, which forms a biclique motif (Figure 1a, bottom). We say that this motif was formed by a wiring rule, expressed by the operator ***O***, which we can write as (101*XX*)***O***(110*XX*). The rule (101*XX*)***O***(110*XX*) should be interpreted as “all nodes that fit the identity 101XX connect (***O***) to all nodes with identity 110XX”. This formalism will become the basis of introducing links into a network for the remainder of the manuscript.

**Figure 1:**
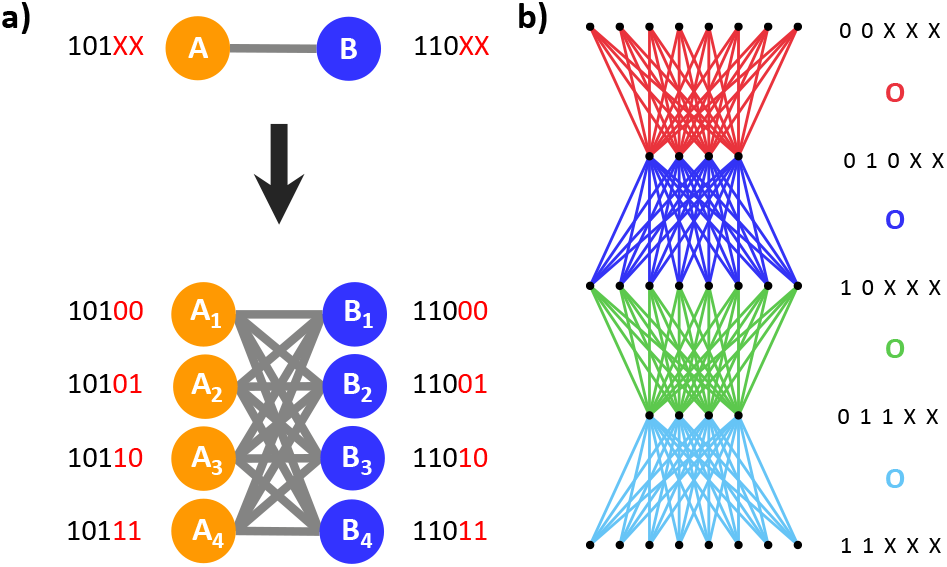
Building structured networks from wiring rules. **(a)** An edge is defined at the set level (top), where sets are uniquely identified by the unique expression of three bits. Each set contains 4 nodes (bottom), each marked by the set-defining bits (black), but also given a unique identity by two additional bits (red). Thus, the single edge between 101 and 110 at the set level can be represented as the rule (101*XX*)***O***(110*XX*) which acts at the node level, producing a total of 16 links. **(b)** Wiring rules generating a 5-layer FFN (left) with 4 rules (right). The generating rules are color-coded, where the colored edges in the network are generated by the colored ***O***s to their right.

The wiring rule of Figure 1a succinctly represents both the number and identities of compatible nodes in a network. In this example bits 1, 2, and 3 were “active”, meaning that their state was crucial for determining which nodes participated. Meanwhile bits 4 and 5 are “passive,” in that they could either be on or off, without including or excluding a node from the rule. If we quantify the number of passive bits as *x* (here *x* = 2), we can see that each set contains 2^*x*^ (here 2^2^ = 4) nodes. Further, if we consider *x*_*S*_ and *x*_*D*_ to be the number of passive genes in the source and destination sets, respectively, then the resulting wiring rule contains 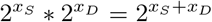 links. In this way, the *active* bits provide *specificity* to wiring rules, while the *passive* bits inform *multiplicity*.

In isolation, a single wiring rule is limited in its application. However, we may choose to impose multiple rules on a system at once. The resulting set of rules can encode a complex topology. To illustrate such representations, we detail two heuristics, one for constructing feed-forward networks, and a second for developing a scale-free network.

We begin with Feed-Forward Networks (FFNs), which have been extensively utilized as a substrate for complex computation in Artificial Intelligence [11], and as a building block, or motif, in complex networks [12]. The simplest FFNs are characterized by a fully-connected layer structure (Figure 1b), such that all nodes in a layer *i* receive connections from all nodes in layer *i−1* and subsequently send connections to all nodes in layer *i+1*. To illustrate this point, we have color-coded links in Figure 1b by the layers that they join. To encode a FFN, we assign nodes in a layer a shared identity. For instance, the 8 nodes in the first layer of Fig 1b correspond to 00XXX, while the 4 nodes in the following layer have identity 010XX. Given this labeling, the connectivity of two adjacent layers will be represented by a single wiring rule, producing directed connections from the “source” set (input to a layer) to the “destination” set (output of a layer). Thus, to link the first layer to the second in Fig 1b, we write the rule (00*XXX*)***O***(010*XX*), which encodes the red edges. In order to have connected layers, the destination set of one rule will be the source set of the next. For Figure 1b, given that the first rule was (00*XXX*)***O***(010*XX*), then the second wiring rule starts with the previous rule’s destination set (010XX), generating the new rule, (010*XX*)***O***(10*XXX*), which now encodes the blue links. If a FFN has 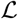 layers, with 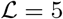 in Fig 1b, then 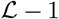 wiring rules are required to encode the network, thus we have 4 unique wiring rules in Fig 1b. In summary, we find that genetic wiring rules allows for an intuitive and succinct generative heuristic for the Feed-Forward Network architecture.

The ease of encoding a FFN can be attributed, in part, to the observation that inter-layer connections form the natural product of wiring rules: bicliques. Yet, genetic rules are capable of generating networks where bipartite motifs aren’t already fundamental units of the topology. To illustrate this, we turn to scale-free (SF) networks, which are characterized by a power law distribution of node degrees [6]. In encoding a SF network, we begin by connecting one node to all nodes; given *b* = 4, we use the rule (0000)***O***(*XXXX*), as in Fig 2a. This single rule produces one node that connects to all 2^*b*^ nodes, giving *p*(2^*b*^) = 2^−*b*^. Next, we introduce a rule that connects 2 nodes, 0000 and 0001, to 1/2 the total nodes: (000*X*)***O***(*XXX*0), for *b* = 4 (Fig 2b). Given that 0000 is already connected to all nodes, we have 0001 with *k* = 2^*b*−1^, thus *p*(2^*b*−1^) = 2^−*b*^. The overlap in the source set has set *p*(2^*b*−1^) = *p*(2^*b*^), yet the power-law begins to appear when introduce the third rule, where we connect 4 nodes to 1/4 of the total nodes, or (00*XX*)***O***(*XX*00) for b = 4 (Fig 2c). Now, this again doesn’t change the degree of 0000 and 0001, but two nodes, 0010 and 0011, have *k* = 2^*b*−2^, thus *p*(2^*b*−2^) = 2^1^ ∗ 2^−*b*^ = 2^1−*b*^. As we continue this process, we connect 8 nodes to 1/8 of the total nodes, with (0*XXX*)***O***(*X*000) for b = 4 (Fig 2d), producing *p*(2^*b*−3^) = 2^2^ ∗ 2^−*b*^ = 2^2−*b*^. Reaching the limit of the example network size, we connect 16 nodes to 1/16 of the total nodes (or 1 nodes in b = 4 through (*XXXX*)***O***(0000), producing *p*(2^*b*−4^) = 2^3^ ∗ 2^−*b*^ = 2^3−*b*^ (Fig 2e). If we utilize all of the rules in concert, we produce a scale-free network (Fig 2f) with in- and out-degree distributions of *p*_*in,out*_(*k*) = 0.5 ∗ *k*^−1^, a scaling that emerges by observation from the probabilities in this paragraph, and is confirmed by simulation (Fig S1). Although 0 < *γ* < 2 has been shown to be non-graphical for unbounded power law degree distributions [13], our deterministic model bounds the maximum degree to the network size, thereby allowing for graphs in this parameter range (slight changes to the heuristic can produce *p*_*in*_(*k*) ~ *k*^−2^ and *p*_*out*_(*k*) ~ *k*^−0.5^, see S1.1.1). The heuristic for SF networks thus begins with a single rule that connects node to all nodes (e.g. (0000)***O***(*XXXX*)) and iteratively adds Xs to the Source set and removes Xs from the Destination set until the Source set becomes all nodes and the Destination set becomes a single node (see SI 1.1.1 for pseudocode). In summary, we have proposed a heuristic that produces a scale-free network with *p*_*in,out*_(*k*) = 0.5 ∗ *k*^−1^ for a system of size 2^*b*^ while utilizing only *b* wiring rules.

**Figure 2:**
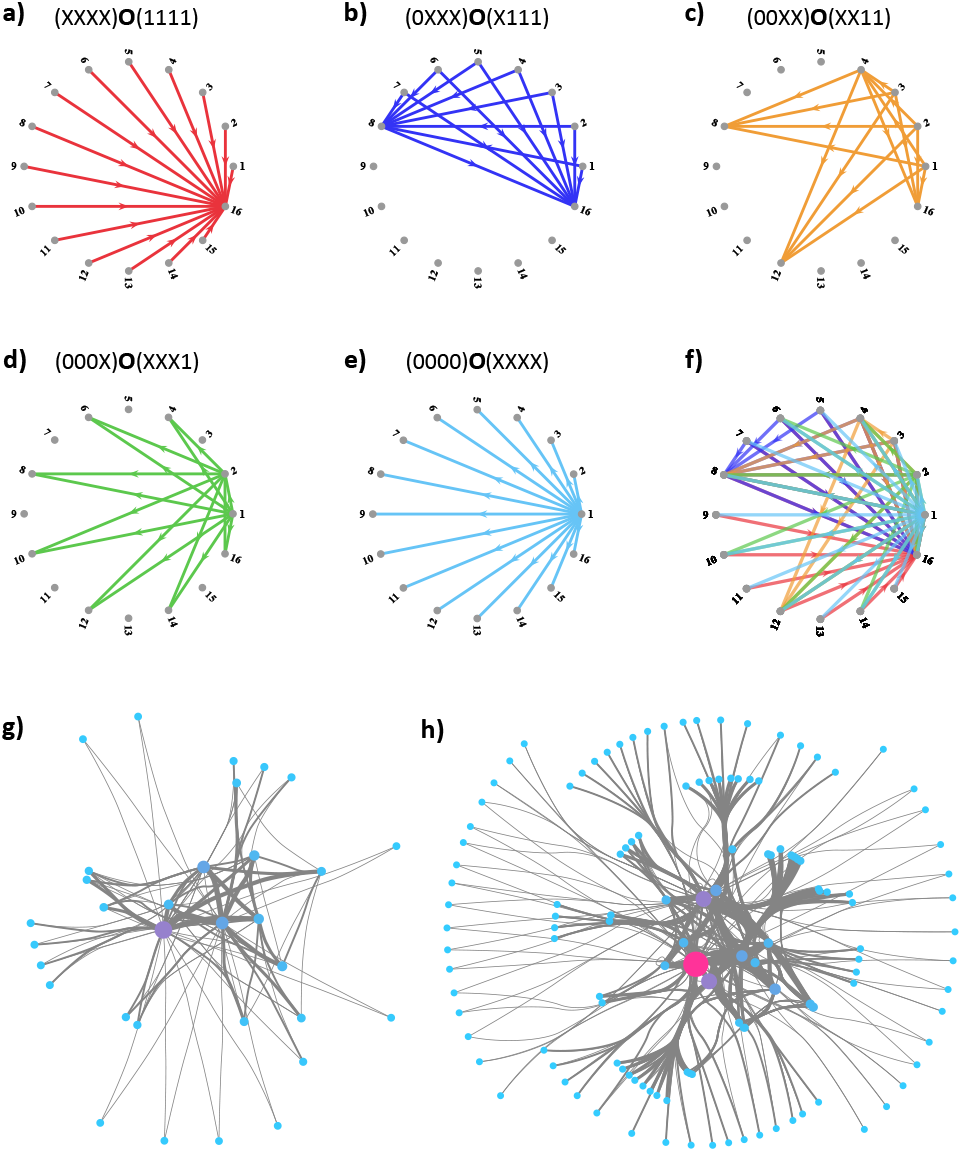
Building scale-free networks using wiring rules. **(a-f)** Wiring rules generating the scale-free graph in (g) with b = 4, where the rule (top) initializes the set of colored edges (bottom). Edges created by each wiring rule are colored in (f), visualizing how individual rules add together to the final network. **(g-h)** Cytoscape visualizations of scale free graphs generated by the model for b = 5 and b = 6, respectively. Node size scales with degree, and the link weights illustrate the number of individual wiring rules coding for the individual edge.

In this section, we have gone from using a wiring rule to generate a single motif, a directed bipartite connectivity called a biclique, to encoding a FFN with 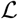 layers using only 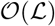 rules (Figure 1b), and a SF network of size 2^*b*^ with 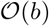 rules (Figure 2). Yet, the proposed modeling framework is not limited to these two case examples, in SI 1.1.2 and 1.1.3 we present two additional succinct heuristics for deterministically generating branching trees and hierarchical networks [14].

### Random Model

The heuristics in the previous section offer a compact representation of known topologies, however the approach does not define a general method for discovering relevant topologies. We therefore propose a random genetic (RG) model that produces a novel family of random networks by probabilistically generating sets of wiring rules.

We begin by observing that the structure of an RG network depends on three variables: *b*, the number of bits determining node identity; *x*, the number of passive bits (number of Xs in a rule); and *r*, the number of wiring rules encoding the network. As before, the value of *b* sets the size of network to *N* = 2^*b*^ nodes. Each of the *r* wiring rules introduces links into the network through a random, but structured, process: on either side of a rule, *x* bits are set to *X* in the rule (are *passive*), and the remaining *b* − *x* bits are randomly sampled from {0, 1} (are *active*). For instance, given *b* = 4, *x* = 2, and *r* = 3, we may introduce the three rules: (*XX*11)***O***(*X*01*X*), (1*X*0*X*)***O***(*X*0*X*1), and (0*X*0*X*)***O***(*XX*01). Note that each of the *r* = 3 rules contains *b* = 4 total bits, of which *x* = 2 are Xs and the remaining bits are chosen randomly. In summary, RG networks contain 2^*b*^ nodes and introduce *r* rules, each of which links two randomly picked sets of nodes, given *x*.

On first pass, if we are given *x* we can infer that 2^*x*^ neurons belong to each set, thus each genetic rule introduces 2^*x*^ ∗ 2^*x*^ links into the network. However, allowing for arbitrary placement of Xs may lead to overlapping sets: two, or more, rules may code for the same links. This is illustrated in Figure 3b, where we introduce the overlapping rules (101*XXXX*)***O***(0*XX*10*XX*), whose edges are marked in green, and (101*XXXX*)***O***(010*XXXX*), whose edges are marked in blue. These rules have the same source set (left of the ***O***), and overlap partially in the destination set (right of the ***O***), leading to a number of edges being coded twice (Figure 3b, red).

**Figure 3:**
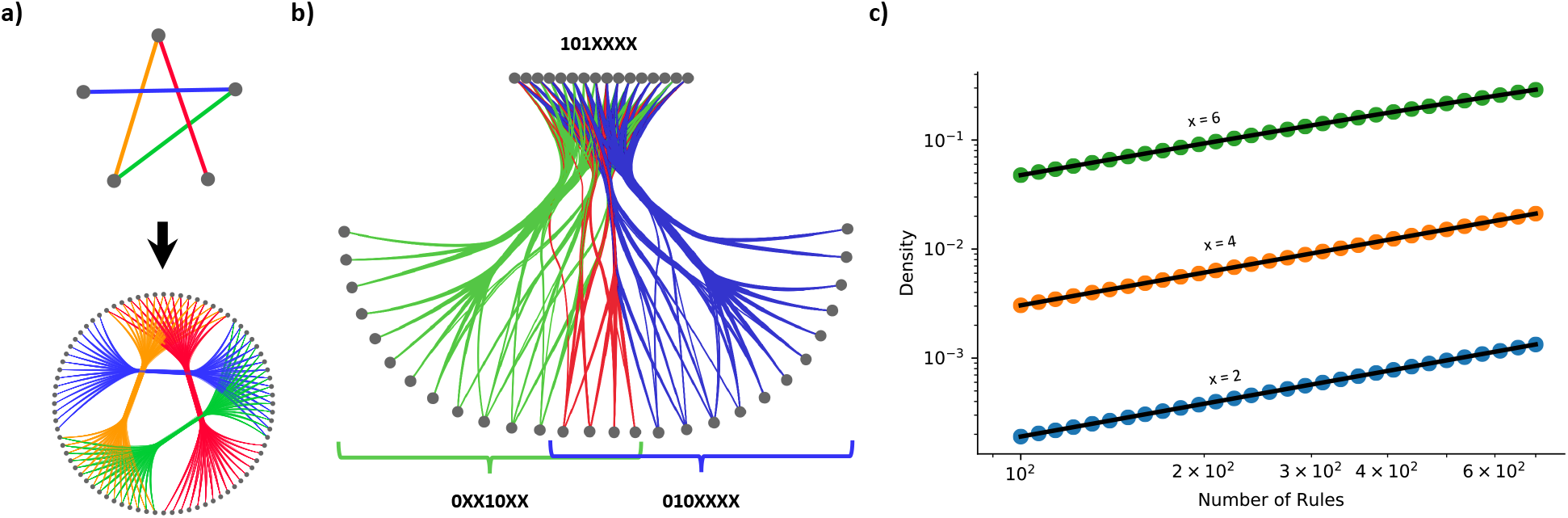
From Wiring Rules to Networks. **(a)** The transition from set level (top) to node level (bottom) can occur not just a single link, as in Figure 1a, but also on a whole network. Here, each set contains 16 nodes, thus each set-level link (top) produces 256 links in total at the neuronal level (bottom). This process of mapping rules from sets to neurons is visualized by link colors. **(b)** Each wiring rule introduced into a network should contribute *L* = 2^2*x*^ links in fixed X model. However, two introduced rules have the possibility to cover the same links if both their source and destination node sets are not disjoint, thus the effective number of links introduced into the network can be lower. Here, two rules are introduced: (101*XXXX*)**O**(0*XX*10*XX*) (green) and (101*XXXX*)**O**(010*XXXX*) (blue), where the left hand side of the rules describes the same nodes: 101XXXX (top). However, the right hand side of the rules also partially overlap, jointly containing 01010XX, leading to the red links in the network being coded twice. Thus, the two rules do not introduce 2^4+4^ + 2^4+4^ = 512 links, but only 2^4+4^ + 2^4+4^ 2^4+2^ = 448. **(c)** Density of RG networks of 4,096 nodes (*b* = 12) with respect to number of rules added. Scatter plots are averaged over 20 simulations, while the black lines are the analytical solution derived in *Random Model*. The fit between simulation and theory holds over multiple Xs, as well as multiple scales of density.

The possibility of rule overlap complicates analytical derivations of network metrics, as considerations for this overlap must be made. For instance, it becomes simpler to calculate the density of the system in terms of the probability of a link not being present, rather than considering the combinatorics of multiple genetic rules coding for one link. The probability that a link is not coded for by a single rule is 1 − *π*^2^, where *π* = 2^*x*−*b*^ is the probability that the rule contains the two nodes the link connects. Thus, the probability that a edge is not covered by *any* rule is (1 − *π*^2^)^2*r*^, where we have two times the number of rules since the nodes can be included on either side of the rule (undirected edges). Consequently, the overall density of such a network can be written as 1 − (1 − *π*^2^)^2*r*^, a prediction that matches numerical simulations (Figure 3c).

Next, we calculate the degree distribution of an RG network via a two-step process: first, we consider the probability *P*(*r*_*i*_) that a node *i* participates in *r*_*i*_ rules. This determines the total possible links it can have: *r*_*i*_ ∗ 2^*x*^. Second, we determine the probability *P* (*k*|*r*_*i*_) that node *i* with *r*_*i*_ rules has a degree *k*. In this way, *P* (*k*) that a given node *i* has degree *k* can be written as

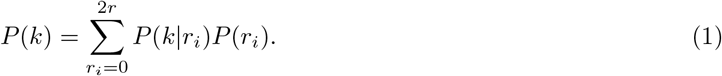

Since each rule connects to node *i* independently, *P*(*r*_*i*_) is a binomial with parameters 2*r* and *π* = 2^*x−b*^ (the probability of node *i* being connected to one specific rule),

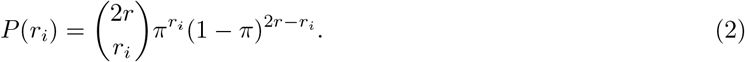

Calculating the second component, *P*(*k*|*r*_*i*_), would require us to take into account all possible overlaps with all possible sizes and their probabilities. We instead continue with an approximation of *P* (*k*|*r*_*i*_), allowing us to illustrate the form of the expected solution.

We consider the naive approximation that the probability *q* that a randomly selected node is *captured by any of the r*_*i*_ *rules node i is connected to*. The probability that a node is connected to *any* of the *r*_*i*_ rules can be approximated as the density of the network given *r*_*i*_ rules:

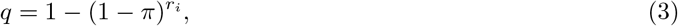

Consequently, the probability that exactly *k* nodes have such a property follows again a binomial distribution,

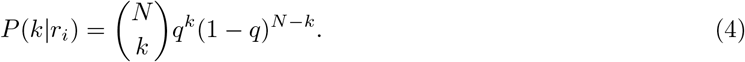

Combining equations (4), (2) and (1), we find that the approximation matches the numerical simulations (Figure 4d-f), although the analytical result overestimates the smoothness of the degree distribution. In other words, our current approximation does not account for the discreteness of overlaps that come from inserting 2^2*x*^ edges with each rule. Intriguingly, the random sampling of the rule sets leads to the overall shape of the degree distribution being binomial (eq. (2)), just like the G(n,p) Erdős-Rényi model [15], and only the wiring rule overlaps produce small deviations from the binomial.

**Figure 4:**
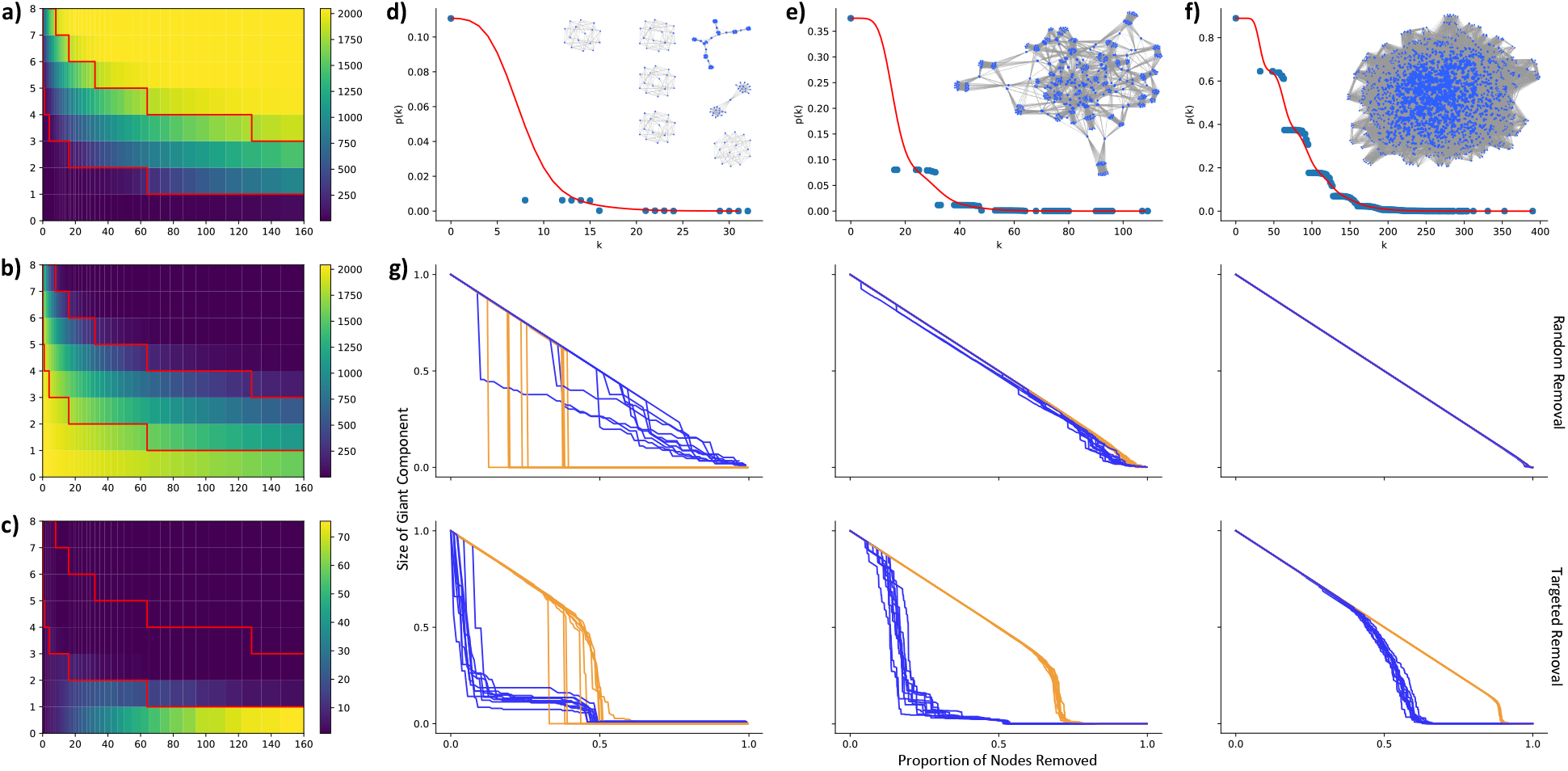
Phase Transition in Random Networks. **(a)** Size of Largest Connected Component (LCC, colorbar) as a function of x (y-axis) and r (x-axis) for a network of 2,048 nodes (b = 11). Red lines indicate the estimated transitions between subcritical to supercritical (lower) and supercritical to connected (top) (see SI 1.3.1 for derivations). The extremes are apparent: below the critical value *N*_*LCC*_ ~ 2^*x*+1^ and above the second line, *N*_*LCC*_ ~ 2^*b*^ = *N*. **(b)** Number of Connected Components (CC) as a function of x and r for a network of 2,048 nodes (b = 11). The extremes are flipped with respect to LCC: below the critical value *CC* ~ 2^*b*^ = *N*, as most nodes do not participate in a wiring rule, thus are their own component, while above the second line, *CC* ~ 1 as all nodes join the LCC. **(c)** An alternative Number of Connected Components (CC) as a function of x and r, where only nodes with *k* > 0 are considered. This biased metric highlights the phase transition near 〈*k*〉 = 1, and reinforces the single CC in the supercritical regime, meaning that none, or at least very few, ‘straggler’ wiring rules are possible outside the LCC above the critical value. **(d)** Degree distribution and example network (inset) from subcritical regime (b = 11, x = 3, r = 15). Blue points indicate the measured degree probabilities over 1,000 numerical simulations, and the red line corresponds to the theoretical prediction from eq 1. Network consist mostly of independent rules, with most nodes participating in no wiring rules (k = 0). **(e)** Cumulative degree distribution and example network (inset) from supercritical regime (b = 11, x = 4, r = 30). This phase consists of a single LCC, although most nodes still participate in no rules, thus have no links (k = 0). **(f)** Degree distribution and example network (inset) from connected regime (b = 11, x = 5, r = 70). Most to all nodes now have links, and therefore participate in the LCC. The ubiquity of wiring rule overlaps in this phase leads to a more consistent fit of the predicted degree distribution (equation 1). **(g)** Robustness of RG networks (blue) and ER networks of equivalent density (orange), under random removal (top row) and targeted attacks (bottom row). Each line corresponds to a single simulation, with 10 initialization plotted for both the RG and th ER network. The 10 initializations are chosen to be characteristic of the overall behavior of the parameter set (additional 100 simulations are shown in Figure S5). Each column’s r and x values correspond to the values set in d-f above. The y-axis measures the relative size of the current LCC compared to the initial giant component, and the x-axis corresponds to the proportion of nodes removed. For both y- and x-axes, the first tick corresponds to and the second tick, where visible, corresponds to 1.

To further characterize RG networks, we next explore the scaling of the Largest Connected Component (LCC) and number of Connected Components (CC). Similar to the Erdős-Rényi random graph model, the RG model displays sub-critical, supercritical, and connected regimes with regards to its LCC [15]. The sub-critical phase is characterized by sparsity and many small clusters, whose number is proportional to the number of nodes in a rule (2^*x*+1^), at which point wiring rule overlap is still improbable (Figure 4d). The phase transition is expected to occur at < *k* >= 1, at which point the network coalesces into a single connected cluster (Figure 4e).

This provides a crucial difference between the ER supercritical phase, which consists of components of size *p*_*s*_ ~ *s*^−3/2^, and the RG supercritical regime, which rapidly coalesces to form a single component. The RG model converges towards a single component as nodes that participated in wiring rules have high *k*, as they received 2^*x*^ links from each rule, thus even though 〈*k*〉 > 1 in the supercritical regime, most nodes will not have any links near the critical point. Therefore, we observe a single dense connected component (Fig 4c) with many high-degree nodes (Fig 3e), while the remainder of the nodes are disconnected. The network is further densified by increasing x or b, thereby reaching the connected phase, where all nodes have *k* > 0 and participate in the LCC (Figure 4f). We estimate phase transition points in SI 1.3.1, where we also illustrate the results should hold independent of network size (i.e. varying b). In summary, we find that RG networks separate into distinct phases at densities similar to ER networks, however this occurs through the formation of a single giant component near the critical point, which then accrues previously unconnected nodes as further wiring rules are introduced.

Although RG networks may seem similar to ER graphs in global behaviors, we turn to network robustness to highlight a key topological difference. The network robustness approach quantifies the degree to which a network stays connected as nodes are removed [16, 17]. We study both random attacks, where nodes are chosen randomly and removed, and targeted attacks, where at each step we remove the current highest degree node. For each type of attack, we ran 100 simulations, both for RG networks (blue) of fixed *r*, *x*, and *b*, as well as on ER graphs of equivalent density (orange), and we quantified the size of the LCC compared to the overall size of the system as nodes were removed (Figure 4g). We find that under random attack, subcritical RG networks have a gradual decay in giant component size, whereas ER networks display an ‘all or nothing’ collapse, as illustrated by the vertical drops under a single removal (Figure 4g top left). However, as the networks densify, the random attacks on RG networks begin to approach the behavior of the ER networks, until the ER and RG networks roughly fall apart at similar rates (Figure 4g top middle, top right). In contrast, RG networks are more vulnerable to targeted attacks (Figure 4g, bottom), especially when fewer rules are introduced. We attribute the differences to the biases of the network encoding: wiring rules introduce dense subgraphs, and removing a single node randomly from a single rule still retains its structure (simply taking it from size SxD to size Sx(D-1), for instance). However, LCCs are formed from the overlap of wiring rules, where a subset of nodes participate in multiple rules, linking the network together. Targeting these overlap hubs leads to a rapid destruction of the LCC, reducing it to the wiring rules it was constructed from. This behavior is conserved by the bottom and left plots of Figure 4g, where the RG networks rapidly decay under targeted attack, but then plateau at 10-20% of initial network size, which corresponds to the relative size of a wiring rule to the giant component. This behavior is markedly different from the one observed for an equivalent ER network, where links are uniformly placed, thus hubs are rare. In summary, we find that while RG and ER networks respond similarity to random attacks in the supercritical and connected regimes, RG networks are much more sensitive to targeted attacks under all regimes.

## Discussion

In this paper, we abstracted a core wiring principle from the brain, where nodes connect based on identity-dependent compatibility, in order to introduce a method of genetic network generation. We show that, when implemented with simple heuristics, such wiring rules can reproducibly construct structured topologies, such as feed-forward, hierarchical, and scale-free networks. We also introduced a Random Genetic (RG) model, where randomly chosen sets of wiring rules allow for the exploration of a novel family of networks. We derive the density and degree distribution of the system, and find that, while the system displays three phrases akin to ER graphs (subcritical, critical, and supercritical link density), RG networks rapidly converges to a single connected component and are sensitive to targeted attacks. In summary, we present a wiring rule based encoding of networks that offers a physically-motivated method for generating and exploring topologies at the boundary of random and structured.

Given its foundations in developmental neuroscience, the proposed network encodings can provide relevant null models for studying the brain. For instance, the mapping of set-level connectivity (top of Figure 1a and 3a) to node-level connectivity (bottom of Figure 1a and 3a) can be considered equivalent to moving between neuronal and cell-type connectivity in neuroscience: each node (neuron) belongs to only one cell type, and connectivity rules are defined between cell types [18, 19, 20]. This can be seen as a RG model with non-overlapping sets, and we derive expected network metrics of the node-level connectivity as a function of set-level connectivity in SI 1.2.

Revisiting the proposed connectivity framework’s biological inspiration can also provide new avenues for exploration. We began by considering how neuronal identity can be modeled as combinatorial expression of *b* cell adhesion molecules, where the wiring rules are interactions of such molecules. Despite the numerosity of cell adhesion molecules for synapse formation, the number involved in a single recognition event is still limited, hence the fixed number of Xs in the RG model [21, 22, 23]. However, actual wiring rules are not expected to utilize the same number of genes each time. This could prompt an alternative model, where the parameter *x* is replaced with *p*(*x*), meaning that each bit in a rule has a probability of being X, rather than *x* bits being set to X every time. Furthermore, in the current model we allowed node identity to be any binary barcode, actual cell identity is though to be sparse, meaning that barcodes would be biased towards containing 0s. Calculations for these two alternative assumptions would require a more explicit treatment of wiring rule overlaps, and therefore would also expand the analytical solutions for the current model.

Finally, the relevance of real-world networks lies not only in their topology, but also the dynamics they support. In this manuscript we catalog network robustness, which proves relevant for neuroscience, as it parallels the brain’s stability as aging, disease, or trauma removes individual neurons. Indeed, random attacks can be seen as modeling neuronal cell death that occurs early in development. Such removals do not impact the integrity of the computational system [24], in fact they are a necessary sparsification step whose disturbance leads to developmental disabilities [25]. On the other hand, targeted attacks, such as head traumas or decline of long-range connections in Alzheimer’s disease [26], have a significant impact in the integrative potential of the brain. We see that the RG networks behave similarly: the topology remains intact under random attacks, but the systems are highly sensitive to targeted removals. Further studies could validate these expected behaviors by simulating neural dynamics on RG networks, thereby gaining a more specific comprehension of the model’s functional capabilities.

## Acknowledgements

We wish to thank Albert-Lészló Barabési for fruitful discussions. D.L.B. was supported by NIH NIGMS T32 GM008313. D.C. acknowledges financial support by the Templeton World Charity Foundation under Grant Number TWCF0268, and by the Hungarian National Research, Development, and Innovation Office under Grant Number GINOP-2.3.2-15-2016-00057.

## Code Availability

We provide MATLAB code for heuristic network generation and python code RG network measurements in the public repository with http://doi.org/10.5281/zenodo.4117624.

## Author Contributions

Both authors have contributed to the design of the project and the writing of the manuscript. Simulations and figures by D.L.B.

## 1 SI

### 1.1 Heuristic Rules

#### 1.1.1 Scale Free Networks

We can summarize the scale-free heuristic from the main text, which generated networks with *p*_*in*_ = *p*_*out*_ ~ *k*^−1^, using the pseudocode:

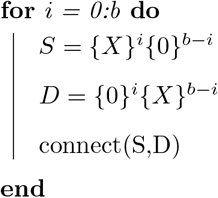

Here “connect” introduces the rule (*S*)**O**(*D*) into the network. For instance, for b = 4 we connect 0000 to XXXX, 000X to XXX0, 00XX to XX00, 0XXX to X000, and XXXX to 0000 (Figure 2, Main Text). The above code creates a directed network with nodes having equal out- and in-degree, while replacing a ‘0’ with a ‘1’ either in the S or D variable results in a network with separate in-hubs and out-hubs.

**Figure 1:**
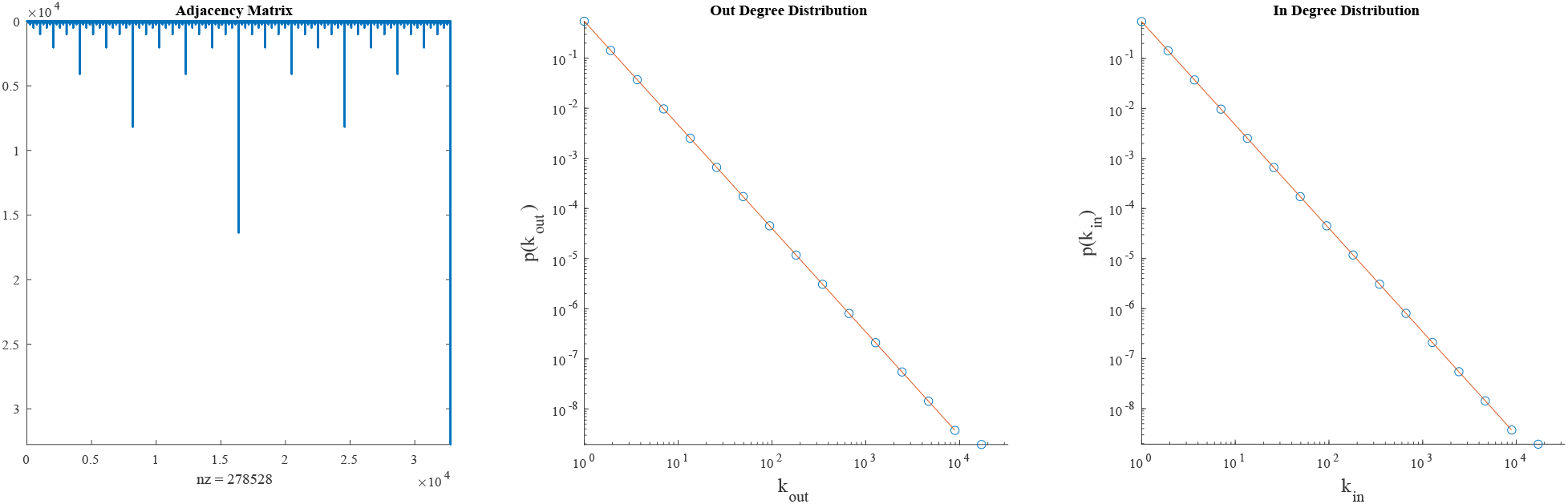
Scale-Free distributions using the connectome model. Left: Adjacency matrix resulting from b = 15. Middle, right: Degree distributions of scale-free graph of b = 15. Both in- and out-degree distributions are best fit by *p*(*k*) = 0.5 ∗ *k*^−1^, as long as the outlier highest degree is excluded.

Through slight modifications of the heuristic, we can produce scale free networks with alternative *γ*. For instance, we can use *b*/2 rules, where we construct D sets as before, but add two Xs to each S set per iteration, resulting in *p*_*in*_ ~ *k*^−2^ and *p*_*out*_ ~ *k*^−0.5^ (Figure S2):

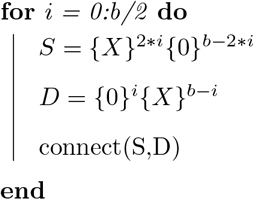

In the case of b = 6, this would produce the rules (000000)**O**(*XXXXXX*), (0000*XX*)**O**(*XXXXX*0), (00*XXXX*)**O**(*XXXX*00), and (*XXXXXX*)**O**(*XXX*000).

**Figure 2:**
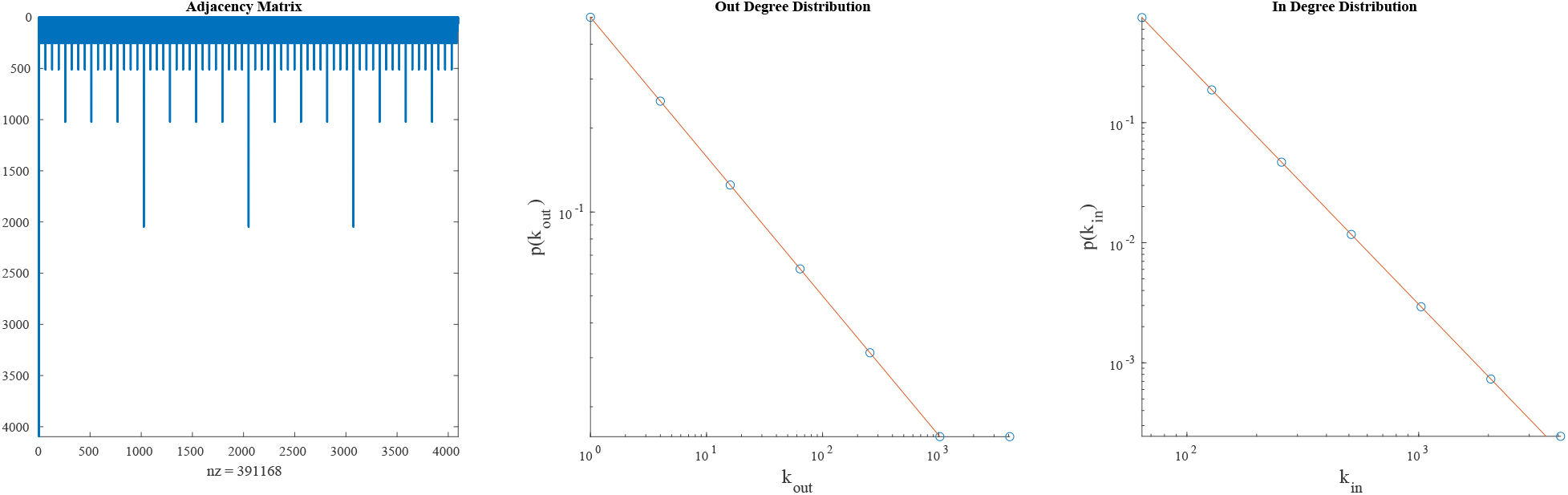
Alternative Scale Free Distribution. Left: Adjacency matrix resulting from b = 12. Middle, right: Degree distributions of scale-free graph of b = 12. While the standard model has matching in- and out-degree distribution, this alternative model has an out-degree distribution is best fit by *p*_*out*_ ~ *k*^−0.5^, and in-degree distribution is best fit by *p*_*in*_ ~ *k*^−2^.

#### 1.1.2 Binary Tree

A simple heuristic for a binary tree of depth *b* can be implemented on {0, 1}^*b*^. The binary barcode of a node can determine its depth in the tree: the location of the first ‘1’ in the barcode (first from the left) defines the depth. For instance, in a tree of total depth 4, 0001 is the root of the tree, and 0100 occurs at depth 3 (Figure S3a). The heuristic consists of a buffer of identities, initialized with the root node. At each step, a barcode is removed from the buffer. A new rule is introduced by using the current barcode as the source set, while the destination set requires removing the first bit of the barcode and adding an X to the end. Then, the two new nodes that are invoked in the destination set are added to the buffer. This process repeats until the destination nodes are selected from the terminal depth (ID starts with ‘1’), at which point the destination nodes are not added to the buffer.

To illustrate, let us again consider a tree with depth b = 4. At initialization, the buffer consists of only the root node, 0001 (Figure S1a, top). We introduce a rule with source set 0001. The destination set is created by dropping the first ‘0’ from the source node, and adding an ‘X’ at the end: 001X. This results in the rule (0001)**O**(001*X*). We then add the two nodes in the destination set to the buffer, which now consists of 0010 and 0011. These will then lead to two new rules: (0010)**O**(010*X*) and (0110)**O**(011*X*). After introducing these new rules the buffer will consist of 0100, 0101, 0110, and 0111. This will lead to the third set of rules: (0100)**O**(100*X*), (0101)**O**(101*X*), (0110)**O**(110*X*), and (0111)**O**(111*X*). However, this step will not place new barcodes in the buffer, as we have reached the final depth (there is 1 as the first bit). Thus, once these rules have been introduced the buffer is empty and the resulting binary tree is now complete (Figure S3a).

**Figure 3:**
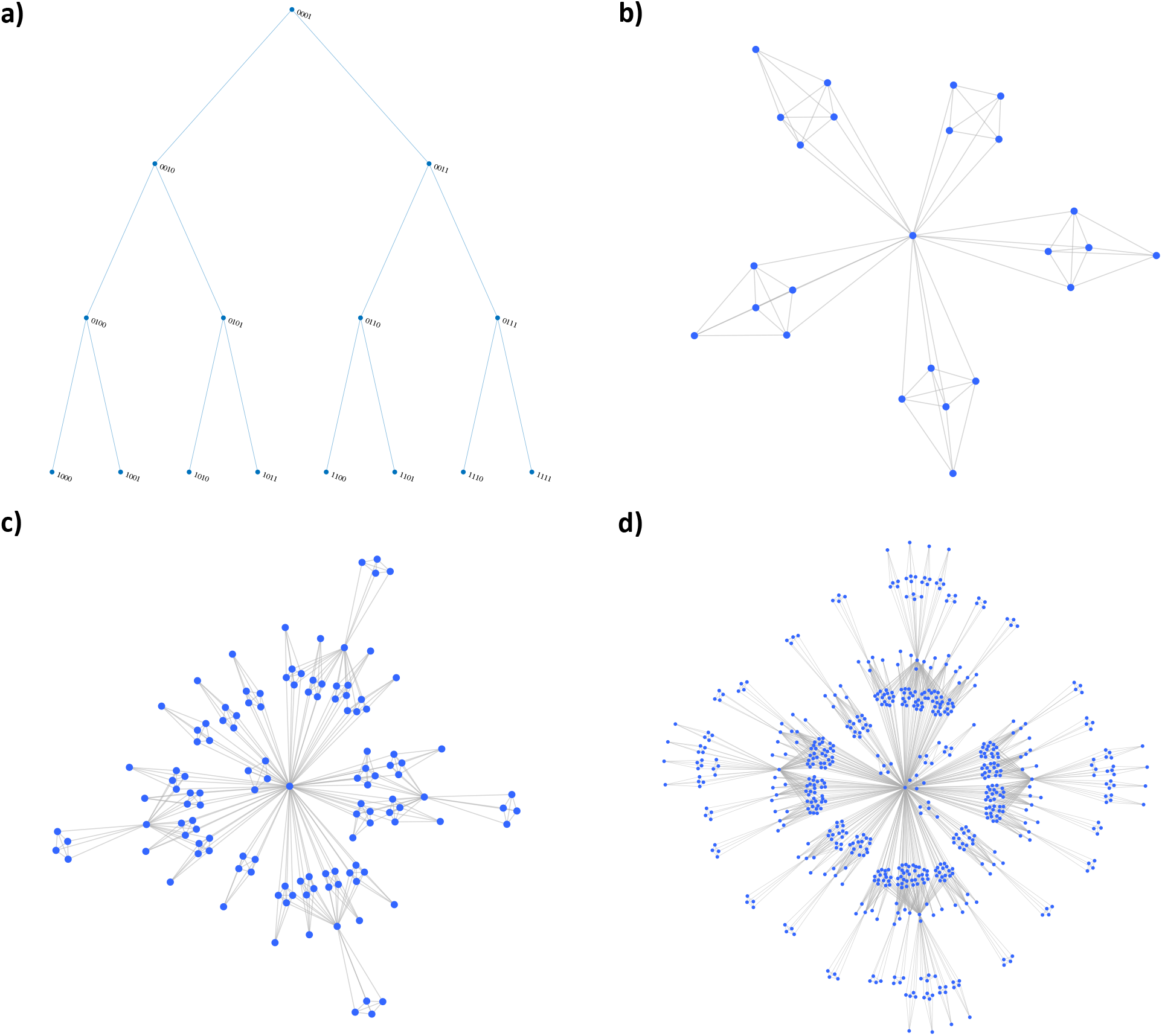
Heuristic Models for Well-Studied Networks. **(a)** Binary tree of depth 4, with node barcodes labeled. **(b-d)** Hierarchical networks for n = 1 (b), n = 2 (c), and n = 3 (d).

#### 1.1.3 Hierarchical Network

Similar to the binary tree, a genetic rule-based heuristic for generating hierarchical networks relies on a motivated indexing of node identity [14]. The core module of a hierarchical network is a 5-clique, corresponding to n = 0, which requires at least b = 3 to encode. We label the node at the center ‘000’, and the four nodes on the ‘periphery’ to be ‘1XX’. In this way, the first bit of a 3-bit set corresponds to whether a node is in the center (0) or at the periphery (1). Thus, the n = 0 hierarchical network can be coded with the rules (1XX)O(1XX), which connects the peripheral nodes to each other, and (1*XX*)**O**(0*XX*), which connects the periphery to the center.

The n = 1 hierarchical network is generated by making four copies of the 5-clique, and having the peripheral nodes of the copy connect to the center node of the original (Figure S3b). This leads to a new definition of core and periphery: a node can be in the ‘core’, or original, clique, or in the peripheral, or copied, cliques, but also a node can be in the center or on the periphery *within* a clique. In this way, n = 1 can be coded easily with 6 bits, the first of which determine which clique a node is in, and the second of which determine where in the clique the node is present. For instance, 101000 is a central node in a peripheral clique, as the first three bits ‘101’ indicate its peripheral clique status, but ‘000’ as the second three bits corresponds to a central node. Alternatively, 000100 is a peripheral node in the central clique. To wire the system up properly, we introduce a separate clique wiring, as introduced in the previous paragraph, for every possible inner and outer clique in the system, such that we have rules of the type (0001*XX*)**O**(0001*XX*), (1001*XX*)**O**(1001*XX*), (1011*XX*)**O**(1011*XX*), (1101*XX*)**O**(1101*XX*), and (1111*XX*)**O**(1111*XX*), as well as similar rules ending in ‘1XX’ on the source and ‘0XX’ on the destination. Then, each of the nodes that are ‘doubly’ peripheral connects back to the ‘doubly’ central node with rule (1*XX*1*XX*)**O**(000000).

This process can be continued, such that we keep introducing Xs into the rules until Xs are present in the ‘outermost’ 3-bit set. This means that for a hierarchical network of depth n, we get 5^*n*−1^ clique-forming rules, 5^*n*−2^ rules that generate an n=1 hierarchy, and 5^*n*−*k*−1^ rules that generate a n = k hierarchy. For instance, for an n = 2 hierarchical network (Figure S3c), we had 5^2−1^ = 5 clique rules and 5^2−2^ = 1 n = 2 rules. This process is best illustrated in the supplementary code, which can create hierarchical networks of any depth, such as n = 3 (Figure S3d), which was not present in the original paper on hierarchical networks [14].

### 1.2 Network Properties under Non-Overlapping Sets

In Figure 1a the active and passive bits were separate: the first three bits were active and the last two were passive in defining both Set A and Set B. If this separation is the same for all rules (all rules are defined by the expression of the first three bits), then each node is contained by only a single set (although sets may be chosen multiple times to participate in a wiring rule). Given a set-level connectivity network (top level of Figure 1a and 3a), we can produce a node-level network by inserting links between nodes if the sets they belong to had a link between them (bottom level of Figure 1a and 3a). We can consider the resulting network to be a function of the original one, such as in SI 1.2.1, we have ER networks at the set level, and *GM*(*ER*) networks at the node level (read *GM*(*ER*) as “node-level Genetic Model of set-level Erdős-Rényi network”). Although this process greatly amplifies the number of nodes and links between the class-level and the neuronal-level networks, we find that the network metrics remain largely unchanged in the process, even if the class sizes are non-uniform (if *x*, or the number of passive bits, varies between cell classes, SI 1.2.2).

#### 1.2.1 Constant Set Size

We previously considered how we can map edges between sets to fully connected groups of participating nodes. Here, we consider how network metrics are impacted by this process. We begin with an ER model at the set level, consisting *L* = *r* links between 2^*b*−*x*^ sets. In this section, each set contains the same number of nodes, 2^*x*^. In this way, each edge between classes corresponds to a wiring rule connecting 2^*x*^ nodes to another 2^*x*^ nodes on the resulting network of 2^*b*^ nodes, which we term *GM*(*ER*). Given *L* = *r* links between 2^*b*−*x*^ sets, at the ER level we have a density of

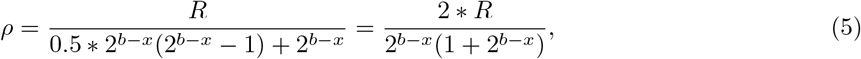

given the sparse assumption of *r* << 2^2(*b*−*x*)^. This is simply the density of an ER graph with *r* links, allowing for self-loops. Performing the *GM*(*ER*) mapping from the set-level ER network to the node-level network with *N* = 2^*b*^ and *L* = *r* ∗ 2^2*x*^, we find that the *GM*(*ER*) has a density of

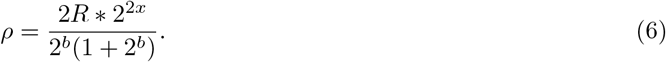

Yet, if we compare the density of the ER and *GM*(*ER*) networks, we find that they are equivalent:

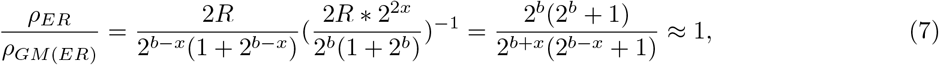

where the approximation holds as long as 2^*b*^ >> 1 and 2^*b*−*x*^ >> 1.

Thus, the mapping of an ER network using constant set sizes does not alter the density of the system. Further, since the clustering coefficient can be defined as the probability that a third edge is present in a triangle, we can assume the clustering will be constant, and *ρ* in the node-level *GM*(*ER*) system as well.

Finally, we derive the degree the distribution of the *GM*(*ER*) system. Since no overlaps between rules occur, the only allowed degree sizes are multiples of the number of links a node gains from a single rule, 2^*x*^. Thus, the degree distribution becomes:

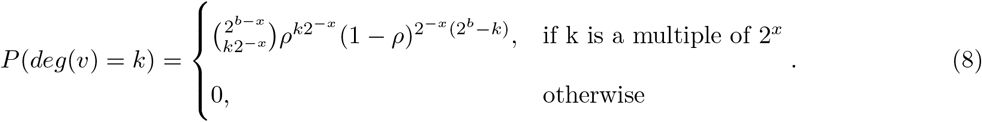

#### 1.2.2 Variable Set Size

In this section we expand the simplifying assumption of mapping networks with constant set sizes, allowing now for sizes to be distributed according to *q*(*s*). The method in the previous section, where we mapped with fixed set sizes, is a special case of this with *q*(*s*) = *δ*(*s* − *s*_0_).

Given an edge between two sets of sizes *s*_*i*_ and *s*_*j*_ drawn from *q*(*s*), *s*_*i*_ ∗ *s*_*j*_ edges are introduced into the node-level network. The expected number of links at the node level is therefore the number of rules multiplied by a factor of 〈*s*_1_*s*_2_〉 = 〈*s*〉^2^. The expected density can then be expressed as

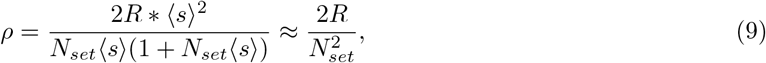

when *N*_*set*_〈*s*〉 >> 1. Thus, the overall density is not expected to differ from the ER network, even with variable set sizes.

To derive the degree distribution, we consider a set with degree *k*_*j*_ at the set-level network. Such a set contains *s*_*j*_ nodes, each with the same degree 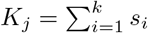. Depending on *q*(*s*) and the degree distribution *p*_*k*_ of the set-level network, the resulting expected degree distribution of the node-level network *p*_*K*_ is given by

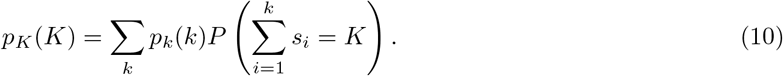

According to the law of total expectation and the law of total variance,

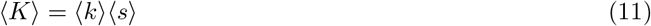

and

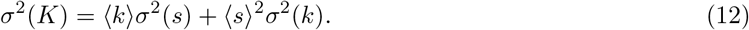

If the set-level model is an ER network, having Poisson degree distribution with *σ*^2^(*k*) = 〈*k*〉, (12) simplifies to 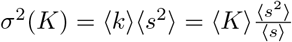. The *GM*(*ER*) model under variable set size therefore has a degree distribution with Fano-factor (as a measure of dispersion) 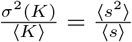.

### 1.3 Behavior of Random Genetic (RG) Model

#### 1.3.1 Critical Points

In this section we propose approximations for the transitions from subcritical to supercritical to connected phases of the RG model. We consider the critical point of the system to be equivalent to an ER graph’s critical point, when 〈*k*〉 = 1. The density of the system was derived in-text, and multiplying by the number of nodes we can write:

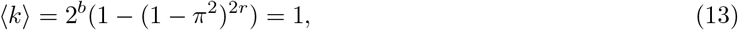

where *π* = 2^*x−b*^. We found it useful to solve for *R*_*crit*_(*x, b*) which is the number of rules introduced to reach the critical point for a given x and b:

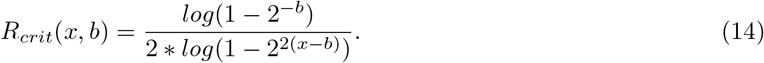

Alternatively, we consider the transition from supercritical to connected regime to occur when the nodes are ‘covered’ by wiring rules, by which we mean that there are sufficient rules for every node to participate in at least 1 rule if there were no overlap. In this manner, *R*_*crit*_(*x, b*) can be found as the number of nodes divided by the number of nodes in a wiring rule:

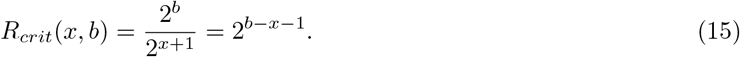

#### 1.3.2 Scaling of Network Metrics

Due to the cost of computation, the in-text results for RG networks were shown for b = 11 (Figure 4, although Figure 3c is b = 12). Introducing many wiring rules and measuring the resulting network properties for larger systems, especially with statistically sufficient repeats, is not computationally feasible. As an indication that our results should be consistent for larger systems, we examine the scaling of density, LCC, and #*CC* with system size (Figure S4). We offer exemplary plots (Figure S4) close to the three *r* values in Figure 4d-f (r = 15, 30, and 70) as well as for r = 160, which is the largest simulated systems in Figure 4a-c.

Although we derived an equation for RG network density in-text, the 1 − (1 − *π*^2^)^2*r*^ formulation does not account for saturation. We thus reframe density as the number of unique links selected from the set of all possible, under replacement. This can be expressed as:

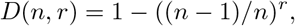

where *n* = 2^*b−x*−1^ ∗ (2^*b−x*^ − 1). We can see that density can be interpreted as a function of *b* − *x*, thus is determined by the relative size of *x* compared to *b*, rather than the system size itself. This observation is confirmed by numerical simulations: the density of the system should follow a similar pattern for larger systems (Figure S4, left column).

The second network metric of interest to the RG model was LCC. We can approximate the size of the LCC by calculating the probability that a node was chosen by a wiring rule. This approximation assumes that a node, if chosen, will belong to the LCC, which we found to be generally valid for the supercritical regime (Figure 4c). We can express the relative size of the LCC as a problem of choosing unique nodes with replacement, allowing us to write:

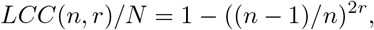

where *n* = 2^*b−x*^. We again find that this is dependent on *b* − *x*, and find that systems of multiple sizes conform to the approximation (Figure S4, middle column). Deviations from the analytical approximation can be attributed to the finite size: in the subcritical regime *LCC* ~ 2^2*x*^, thus *LCC/N* ~ 2^2*x−b*^, from which *x* − *b* is no longer cleanly separable.

The final network metric considered was the number of connected components (#*CC*). This can be seen as the inverse of the LCC: #*CC* is the number of nodes *not* selected by a rule. Thus, we express the relative #*CC* as:

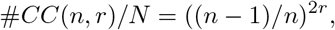

where *n* = 2^*b−x*^ (Figure S4, right column). This again can be parameterized in terms of b-x, except in the connected regime, where #*CC* → 1, thus #*CC/N* → 2^−*b*^. Yet, if we think of LCC and #*CC* as complementary measures, we then have approximates for all three documented regimes.

**Figure 4:**
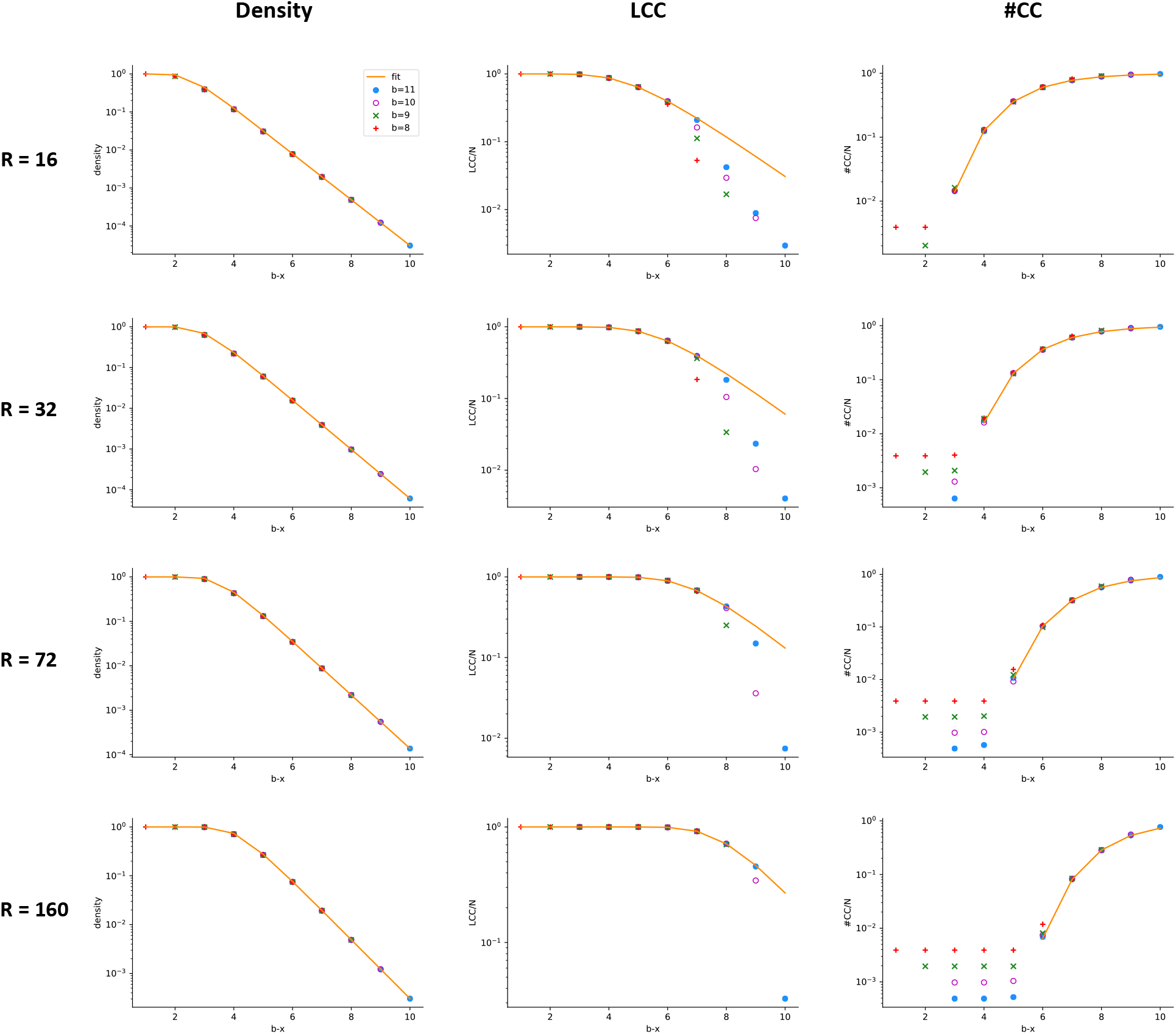
Scaling of Network Metrics with System Size. Plots of Density, LCC and #*CC* are shown for four *r* values. In all metrics and *r* values, the various system sizes display nearly identical scaling, with deviations attributable to finite size arguments, as expressed in SI 1.3.2.

#### 1.3.2 Scaling under Constant Average Degree

An important limit considered in graph models is the scaling of network metrics with system size, when a constant average degree is maintained. Specifically, how does the number of rules *r*(*N*) scale with system size *N* = 2^*b*^ given a hand-imposed scaling of the average degree 〈*k*〉?

Solving for the number of rules given the network density *ρ* = 1 − (1 − 2^2(*x−b*)^)^2*r*^ and average degree 〈*k*〉 = *ρN* gives

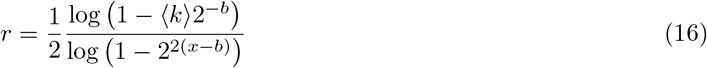

In the limit of *b* >> 1, using log(1 + *a*) ≈ *a*, (16) is simplified to

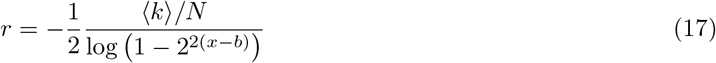

If, in addition, *b* − *x* = *g* is constant, then the number of rules scales as

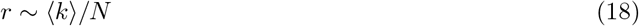

In this limit, therefore, the number of rules scales inversely with system size. This can be understood based on the fact that wiring rule sizes, in terms of the number of new edges they introduce, grow proportionally to the square of the system size *N*.

If, on the other hand, *b* − *x* >> 1 (such as when *x/b* or *x* are kept constant), then

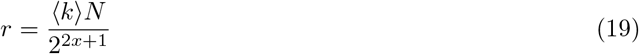

In this regime, the number of rules always scales as the system size *N*.

**Figure 5:**
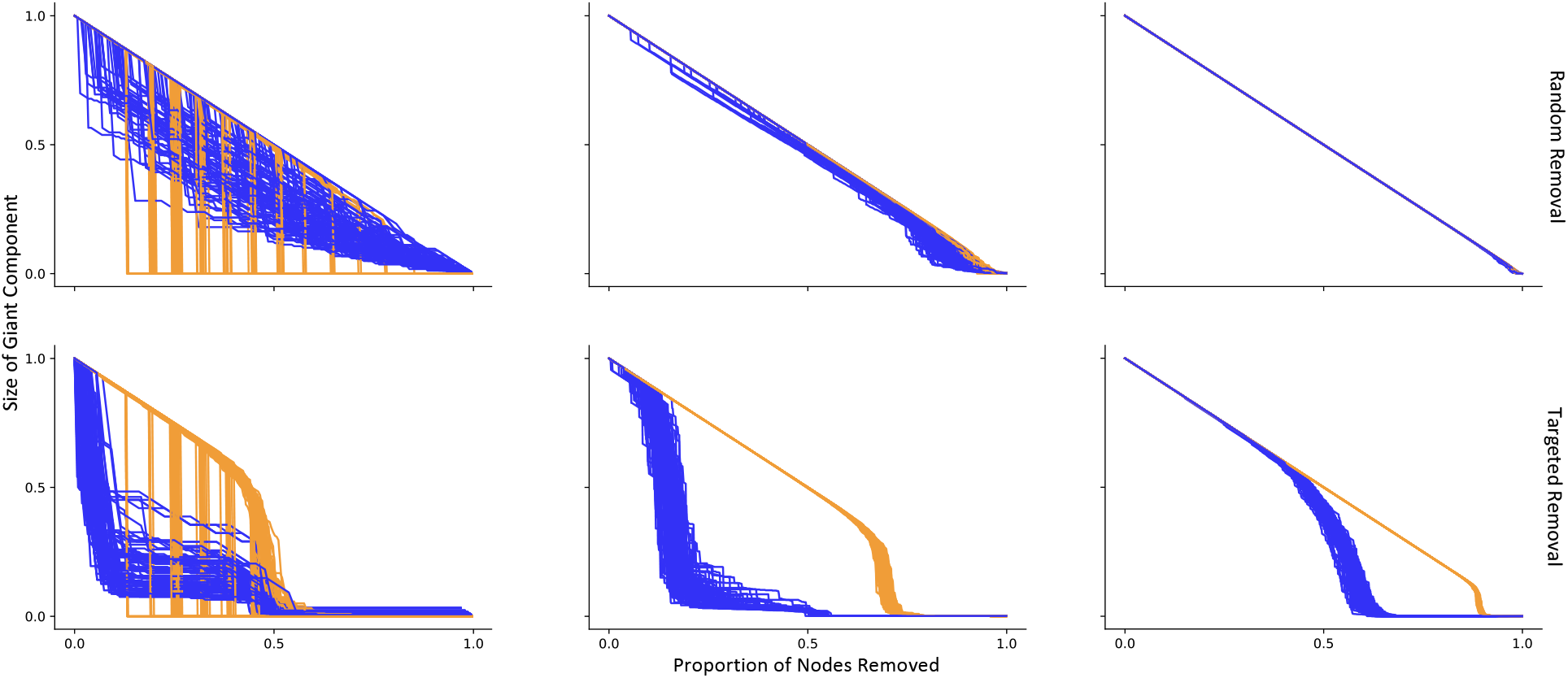
Additional Simulations for Network Robustness. Robustness of RG networks (blue) and ER networks of equivalent density (orange), under random removal (top row) and targeted attacks (bottom row). Each line corresponds to a single simulation, for a total of 100 initialization of each RG and ER parameter set. Each column’s r and x values correspond to the values set in Figure 4d-f above (left: *x* = 3, *r* = 15, middle: *x* = 4, *r* = 30, right: *x* = 5, *r* = 70). The y-axis measures the relative size of the current LCC compared to the initial giant component, and the x-axis corresponds to the proportion of nodes removed. For both y- and x-axes, the first tick corresponds to 0.5 and the second tick, where visible, corresponds to 1.

